# Cryo-EM Structure of the R388 plasmid conjugative pilus reveals a helical polymer characterised by an unusual pilin/phospholipid binary complex

**DOI:** 10.1101/2024.03.04.583355

**Authors:** Abhinav K. Vadakkepat, Songlin Xue, Adam Redzej, Terry K. Smith, Brian Ho, Gabriel Waksman

## Abstract

Bacterial conjugation is a process by which DNA is transferred unidirectionally from a donor cell to a recipient cell. It is the main means by which antibiotics resistance genes spread among bacterial populations. It is crucially dependent upon the elaboration of an extracellular appendage, termed “pilus”, by a large double-membrane spanning secretion system termed conjugative “type IV secretion system”. Here we present the structure of the conjugative pilus encoded by the R388 plasmid. We demonstrate that, as opposed to all conjugative pili produced so far for cryo-EM structure determination, that encoded by the R388 plasmid is greatly stimulated by the presence of recipient cells. Comparison of its cryo-EM structure with existing conjugative pilus structures highlights a number of important differences between the R388 pilus structure and that of its homologues, the most prominent being the highly distinctive conformation of its bound lipid.

## Introduction

Bacterial conjugation is the process by which DNA (usually plasmids or other genetic mobile elements) is transferred unidirectionally from a donor to a recipient cell (Lederberg and Tatum, 1946). It plays a crucial role in horizontal gene transfer, the major means by which bacteria evolve and adapt to their environment. It is also a process of immense biomedical importance since conjugation is the main vector of propagation of antibiotic resistance genes among bacterial populations (Barlow, 2009; Virolle et al., 2020).

In Gram-negative bacteria, conjugation is orchestrated by three large complexes assembling in the donor cell: a DNA-processing machinery known as “the relaxosome”, a membrane-embedded transport machinery termed “type 4 secretion (T4S) system”, and a pilus (Waksman, 2019). Nothing is known about the complexes formed to facilitate transport through the recipient cell membrane.

The T4S system is a multi-mega Dalton secretion apparatus embedded in the double membrane of Gram-negative bacteria. It is minimally composed of 12 proteins termed “VirB1-11 and VirD4” (Chandran Darbari and Waksman, 2015; Costa et al., 2020). Three components, VirB7, VirB9, and parts of VirB10, form the so-called outer membrane core complex (OMCC). The OMCC connects to an inner-membrane complex (IMC) composed of VirD4, VirB4, VirB3, parts of VirB6, VirB8 and VirB10. OMCC and IMC are connected through a Stalk made of VirB5 and parts of VirB6. A periplasmic ring called the Arches, made of a part of VirB8, surrounds the base of the stalk (Costa et al., 2023; Macé et al., 2022). At least two ATPases (VirB4 and VirD4), sometimes three (VirB4, VirD4, and VirB11) power the system.

The conjugative pilus is an essential element in conjugation. It is made of a major component, VirB2, and a minor one, VirB5. Pili may serve either as devices mediating attachment of the donor cell to the recipient cell, or as a conduit for relaxase/ssDNA transport, or both (Beltran et al., 2023; Goldlust et al., 2023). Some conjugative pili are capable of retraction, which will bring donor and recipient cells together. Indeed, tight conjugative junctions have been observed which have led to the suggestion that cell-to-cell contacts are required for conjugation to take place (Low et al., 2022).

The first conjugative pilus structures ever determined, that encoded by the F-family plasmids pED208 and pOX38, demonstrated that conjugative pili are polymers, the unit of which is made of a binary complex of VirB2 bound to a phospholipid that assembles into a 5-start helical filament (Costa et al., 2016). Since then, all conjugative pilus structures from eubacteria that have been solved have confirmed this general architecture, showing variations only in the type and relative positioning of the bound phospholipid and in helical parameters (Amro et al., 2023; Kreida et al., 2023; Zheng et al., 2020).

Recently, we presented the cryo-EM structure of a multi-mega Dalton complex of a conjugative T4S system, encoded by the R388 plasmid (Macé et al., 2022). This study not only delineated how different proteins come together to form a T4S system but also provided novel insights into a potential mechanism for pilus biogenesis. While results of a site-directed mutagenesis study supported the proposed mechanism (Macé et al., 2022), a more comprehensive exploration of this mechanism necessitates a detailed structural analysis of the conjugative pilus encoded by the R388 plasmid. Moreover, any further dissection of substrate-transfer and translocation mechanisms employed by the T4S system to execute conjugation mandates an experimentally derived pilus structure. This structural information is essential for designing targeted mutations capable of trapping the T4S system caught in the act of substrate transport.

In this manuscript, we describe the cryo-EM structure of the pilus elaborated by the R388 T4S system. To our surprise, although the polymerising unit is a binary complex of phospholipid-bound pilus subunit, the lipid is observed in a very different conformation, never observed in conjugative pili, but reminiscent of lipid bound to lipid binding proteins such as lipases and transferases. In effect, while in other conjugative pili, the lipid adopts a configuration of their two acyl chains similar to what is observed in membranes i.e. running parallel to one another, that of the R388 pilus is splayed. Thus, during pilus biogenesis, the lipid extracted from the membrane must undergo a conformational change from membrane-inserted to pilus-bound that is likely energetically costly to stabilise. It is predicted that such pili, once assembled, might be unable to retract.

## Results and discussion

### Pilus production and structure determination

Pili were initially produced and purified according to the previously-described protocol for F and pED208 pili (Costa et al., 2016). However, the yield was very poor. Interestingly, it was observed that in the presence of recipient cells, there was a marked (10-to 20-fold) increase in pilus production by donor cells. Consequently, a co-culture of donor and recipient cells was prepared, and the mixture was plated on solid agar—an essential step for mating in the context of the R388 plasmid. After an incubation period of one hour to facilitate conjugation, the resulting plate was scraped to collect cells. Subsequent purification procedures followed minor modifications to the conventional protocols, resulting in high yields of purified pili. The preparation of EM grids and cryo-EM image processing of the filaments to generate a high-resolution reconstruction of the R388 pilus using helical image processing pipeline implemented in CRYOSPARC are described in Methods (Figure 1 and Table S1). Since a phospholipid molecule was observed bound to each VirB2 pilus subunits (also called TrwL) in the final electron density map, efforts were directed towards identifying this lipid through mass spectrometry, as outlined in Costa et al. (2016) (Costa et al., 2016) and described in Methods (Figure 2). The prominent bound phospholipids were identified as phosphatidylglycerol PG 32:1 and PG 34:1.

**Figure 1.**
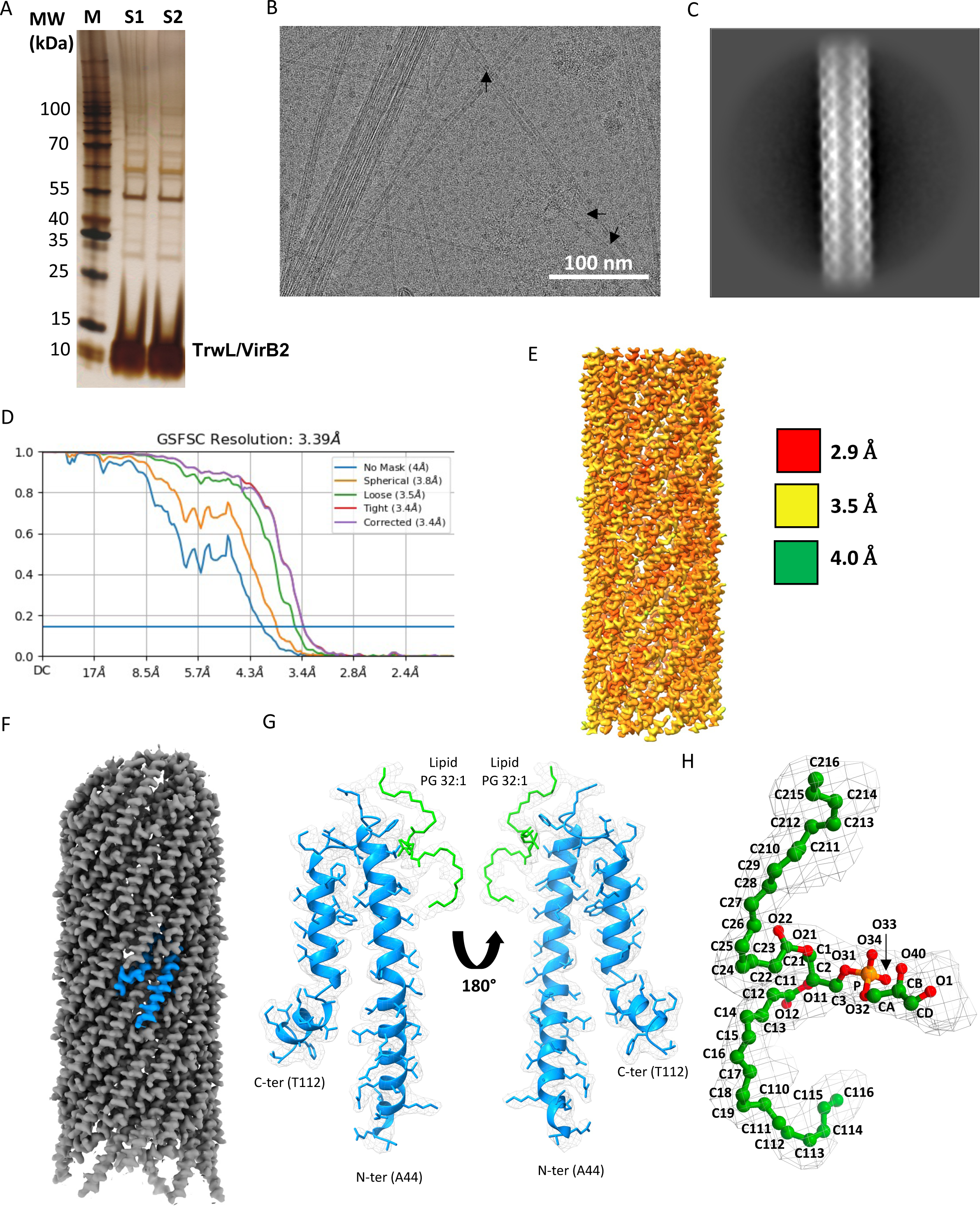
Biochemistry and cryo-EM of R388 pili. (A). SDS-PAGE analysis of the purified R388 pili. Molecular weight markers are indicated on the left. The VirB2 (also called TrwL in the R388 naming nomenclature) band is indicated. Bands were revealed using silver staining. (B). Cryo-EM micrograph of the R388 pili. Arrows indicate examples of pilli. 4884 such micrographs were collected in this work. (C). A typical 2D class of the R388 pilus fibre. (D). Average resolution derived from Fourier Shell Correlation. This FSC plot shows curves for correlation between 2 independently refined half-maps with no mask (blue), spherical mask (green), loose mask (red), tight mask (cyan) and corrected (purple). Cut-off 0.143 (blue line) was used for resolution estimation. (E). Local resolution calculated using CRYOSPARC (FSC cut-off 0.5) and coloured as indicated in the scale on the right of the map. (F). Electron density map of the pilus (sharpened in CRYOSPARC and rendered at a contour level of 0.32 sigma in ChimeraX). (G). Model derived from electron density. The VirB2 pilus subunit is shown in blue ribbon while the PG 32 is shown in stick representation colour-coded in green. Density is shown in chicken wire contoured at 0.25 sigma. Left and right panels show two views distant by 180°. (H). Details of density and model of PG 32 showing the naming nomenclature for all atoms as in Marsh (2003) (Marsh, 2003). The corresponding density is contoured at 0.15 sigma.

**Figure 2.**
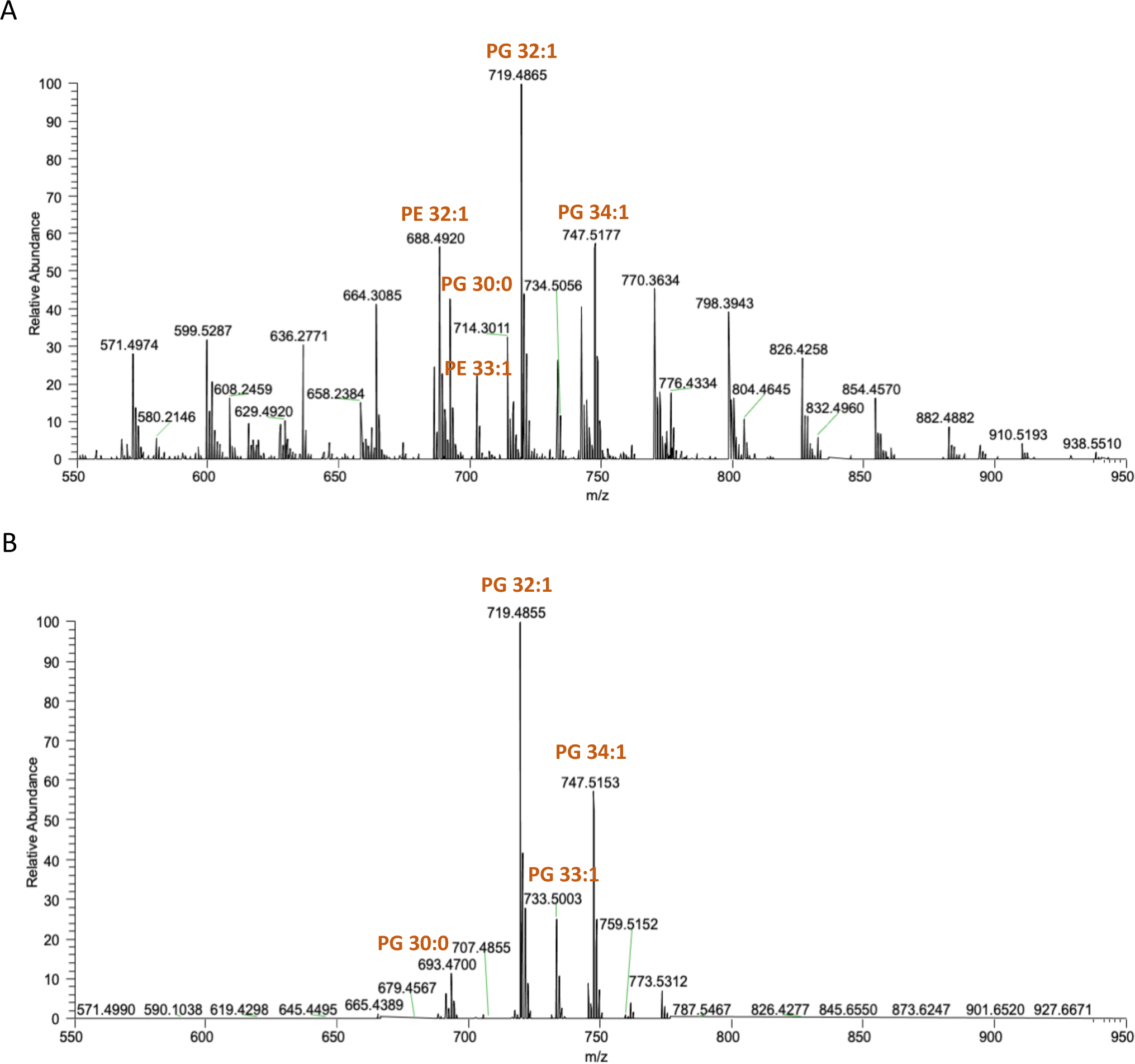
MS analysis of the lipids extracted from pED208 pili. (A) Negative ion mode survey scan (600-780 m/z) of lipid extracts from whole cell membranes. (B) Negative ion mode survey scan (600-780 m/z) of lipid extracts from purified pili pre-treated with PLA2. In all cases, phospholipids identity was confirmed by accurate mass and where appropriate daughter fragmentation.

### Structure of the pilus subunit

The sequences of VirB2 family proteins are overall conserved (Figure 3A) and thus, it is not surprising that their structures should also be conserved. As observed in all pilus structures, the VirB2 pilus subunit is mostly α-helical. In the first structure of a conjugative pilus ever published, that of the F and pED208 pili (Costa et al., 2016), we observed 3 α-helices, which we annotated α1 to α3. α1 and α2 run parallel to α3, forming a helix-loop-helix hairpin between α2 and α3. The loop is positively charged and facing the lumen, while the N- and C-termini remain solvent-exposed. A small unstructured insertion is observed between α1 and α2, but these two helices run along the same axis. Subsequent structures of all F-family pili (Figure 3B) showed the same arrangement. However, the subsequent determination of the structure of the T pilus (Kreida et al., 2023) and that of the pKM101 pilus (Amro et al., 2023), although demonstrating a similar α-helical organisation, showed significant differences, which are also observed in the structure of the R388 pilus presented here. While, in F-family pili, α1 and α2 are distinct helices, in pKM101/T/R388, they are fused. Similarly, while α3 is a single helix in F-family pili, it is split in pKM101/T/R388. Consequently, two distinct families of pilus subunit structures emerge: the F-family subunits (F/pED208/pKpQIL in Figure 3B) and the pKM101/T/R388 family (Figure 3B). In order to account for their similarities and differences, we suggest the following nomenclature for secondary structures of the pKM101/T/R388 pilus subunit: “α1/2” for the fused α1 and α2 helices, and “α3_1_ and α3_2_” for the two helices of the split α3 (Figures 3A-C). Superposition of pED208 against R388 VirB2s, as well as between pKM101/T family VirB2s shown in Figure 3C provides a visual and quantitative account of similarities and differences between these two structural families.

**Figure 3.**
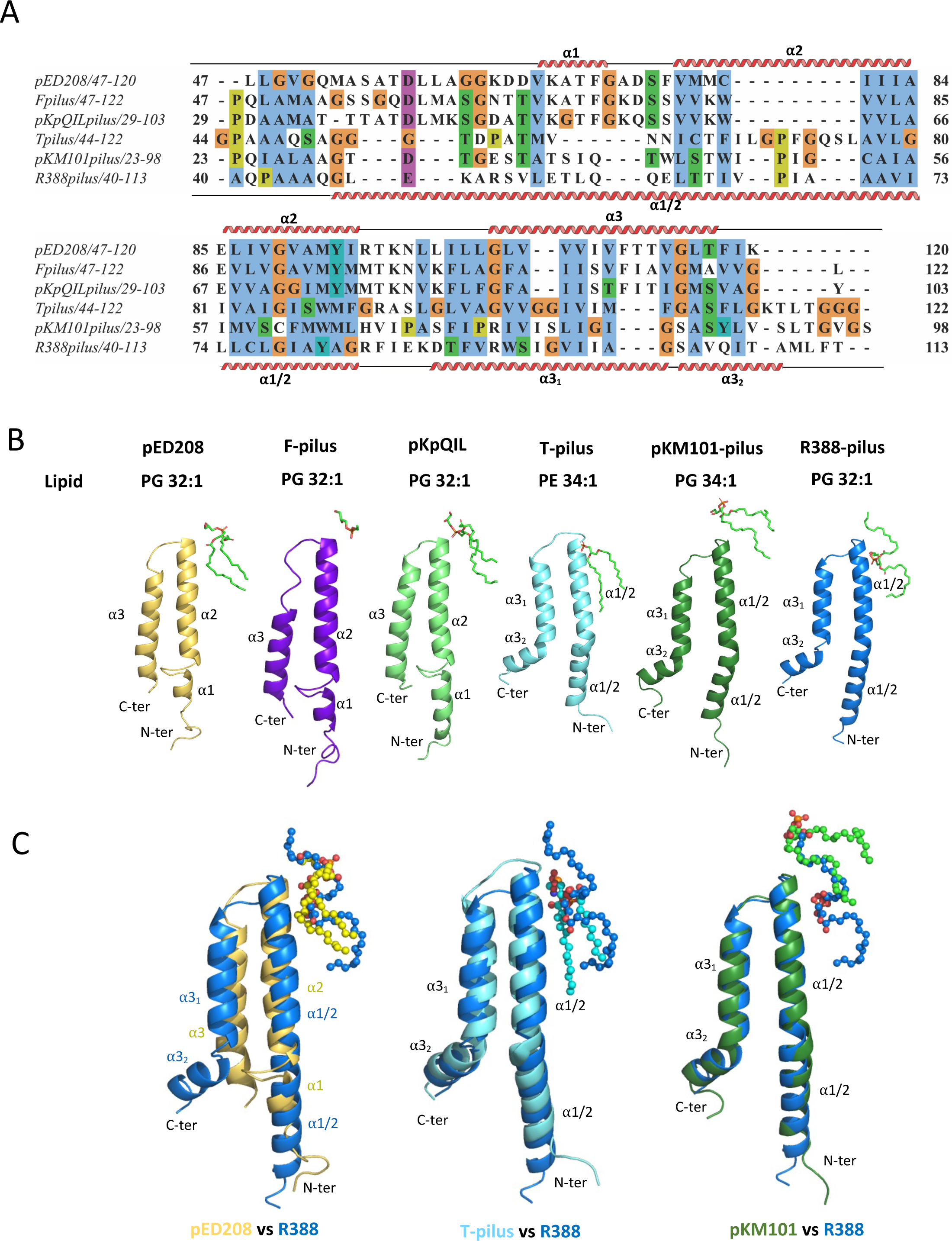
Comparison of sequences and structures of various conjugative VirB2 subunits and their cognate phospholipids. (A). Sequence alignment of VirB2 subunits for which the structure is known. Secondary structures of pED208 VirB2 and R388 VirB2 are shown above and beneath the alignment, respectively, and are named as explained in main text. Colour coding of boxes around residues is as follows: unconserved residues are shown with a white background while conserved hydrophobic residues are shown in blue, negatively charged residues in magenta, polar residues in green, glycines in orange, prolines in yellow and aromatic residues in cyan colour. (B). Structures of the 6 eubacterial conjugative VirB2 subunits for which the structure is known, including the R388 structure from this work. Naming of secondary structures are as in main text. The name of the plasmid encoding the shown proteins is indicated on top of the structure with the type of phospholipid observed bound to these proteins shown just under the plasmid name. PG: phosphatidylglycerol. PE: phosphatidylethanolamine. (C). Superposition of the R388 VirB2-phospholipid unit structure with that of pED208 (left), T pilus (middle) and pKM101 (right). Proteins are in ribbon representation while phospholipids are in ball-and-stick representation. Colour coding of each subunit is indicated at the bottom of each superpositions. The overall RSMD in Ca (in Å) between R388 and pED208, T- and pKM101 are 2.21, 1.23 and 1.1, respectively.

### Structure of the phospholipid

The remarkable observation that all conjugative pili consist of stoichiometric VirB2-lipid complexes was initially made in the context of pED208 and F pili (Costa et al., 2016). In these structures, a density resembling that of a lipid adjacent to each VirB2 subunit was identified. This finding was confirmed in experiments in which the purified pED208 pili were first treated with phospholipase 2 (PLA2) and the remaining bound lipids subsequently extracted and analyzed by mass spectrometry (MS). Two main species bound to the pilin were identified by daughter ion fragmentation as phosphatidylglycerol (PG) species, PG 32:1(16:0, 16:1) and PG 34:1 (16:0, 18:1). These species are also the predominant PG species in whole-cell membranes. However, there was selectivity observed, as there was an absence of the other two major phospholipid classes, phosphatidylethanolamine (PE) and cardiolipin, in the PLA2-treated pili extracts. Moreover, while the total PG pool only accounts for 19% of the total phospholipid content of the *E. coli* membrane, the two major PG species identified in the pilus account for 72% of the lipid content of the pilus.

Subsequent to these initial findings, similar observations on other conjugative pili have been documented, albeit with variations encapsulated within a common theme (Amro et al., 2023; Kreida et al., 2023; Zheng et al., 2020). In one of the most closely related homologs to the R388 system, the pKM101 pilus was noted to exhibit phosphatidylethanolamine (PE) binding to the VirB2 monomer (Amro et al., 2023). In the presented R388 pilus structure, we identify the same lipid composition as observed in pED208 (Figure 2). However, we also observe a strikingly different conformation of the acyl chains for the phospholipid compared to all other conjugative pilus structures (Figure 4A and superposition in Figure 4B). While in pED208 and in all other pili, the two acyl chains, *sn*-1 and *sn*-2, making up the phospholipid run parallel to each other, in R388, the acyl chains are splayed from atoms C13-C15 (atoms 3 to 5 of *sn*-1; Figure S1) and C25-C28 (atoms 5-8 from *sn*-2; Figure S1) with significant differences in the β5-β7 and γ5-γ8 dihedral angles between the pED208 and R388 conformations (Figure S1 and Table S2 (Marsh, 2003)). There are other differences within *sn*-1 and *sn*-2 chains such as in β10 and β11 of *sn*-2 and γ11-13 of *sn*-1, and also in the head group (Figure S1 and Table S2) but they appear less consequential than those responsible for the splaying of the acyl chains.

**Figure 4.**
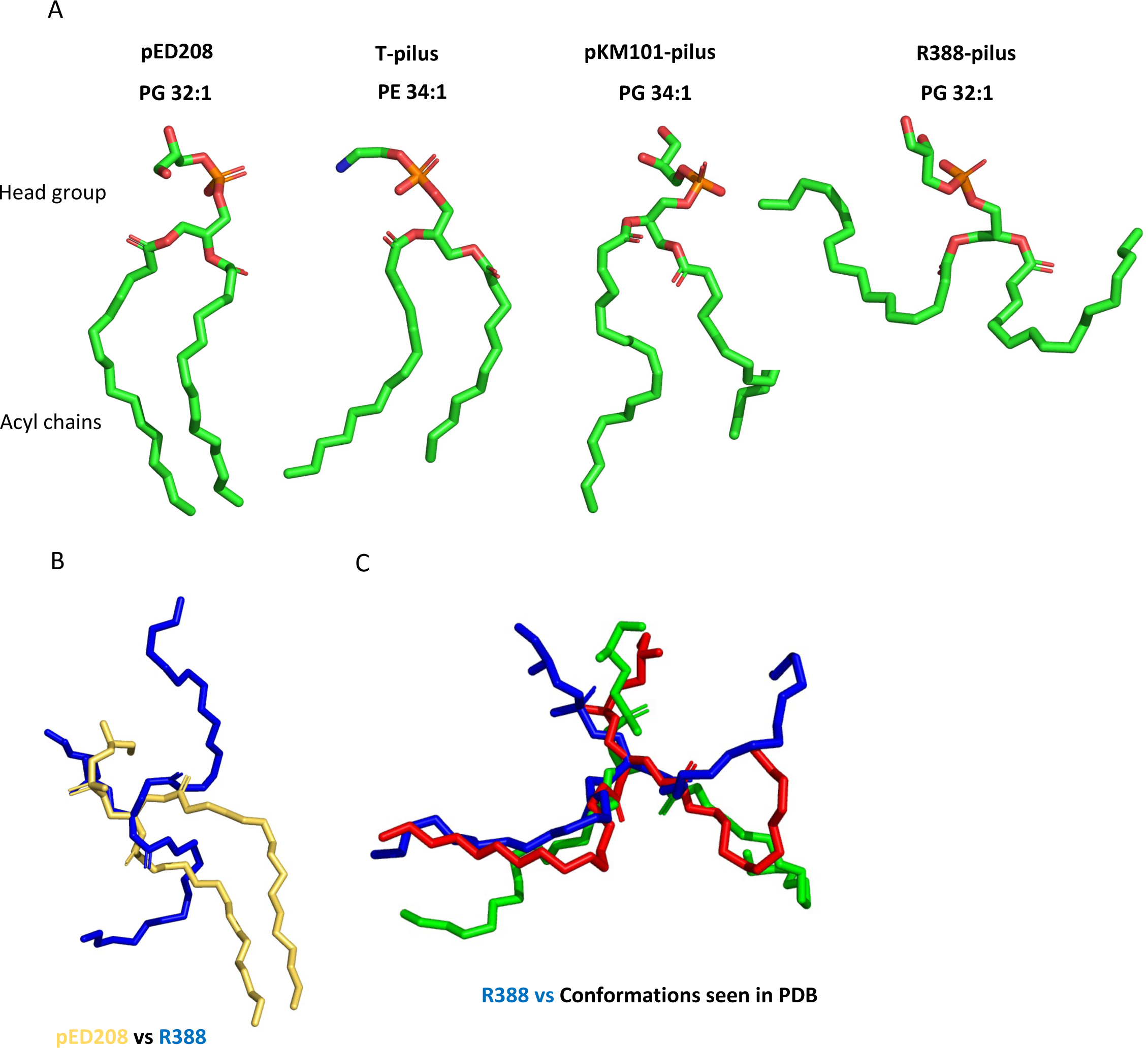
Comparison of the structure of the R388 phospholipid with the phospholipid observed in structures of protein-lipid complexes. (A). Conformation of the phospholipids observed in the structures of the pED208 pilus, the T-pilus, the pKM101 pilus, and the R388 pilus (this work). While the acyl *sn*-1 and *sn*-2 chains in the phospholipids observed in all other pili except R388 run parallel to each other, those of R388 are splayed. (B). Superposition of the structures of the pED208 and R388 PG 32s based on aligning as best as possible the head groups of each structure. (C). Comparison of the R388 (in blue sticks) phospholipid structure with that of other phospholipids bound to 3A0B (in green sticks) or 6LY5 (in blue sticks).

Splayed conformations for phospholipids are known to be energetically unfavourable (Marsh, 2003). In the cellular membrane, the acyl chains of phospholipids run parallel to each other, and therefore to transition to a splayed configuration, the lipid must undergo a conformational change, also likely to be energetically costly. Nevertheless, the splayed conformation is observed, presumably because surrounding subunits form a binding site that stabilises it (see below “protein-lipid interaction” section for details). Splayed conformations have been observed previously in some lipid-interacting lipases and transferases (see examples in Figure 4C). In the case of pili, it might be that such a conformation is rare because many of these pili are retractable, i.e. they are able to depolymerise in a mechanism that involves the reintegration of the subunit-lipid complex within the inner membrane, a process potentially made difficult in R388 by the return of the pilus-bound splayed lipid conformation to the membrane-bound parallel conformation.

### Overall helical assembly

The pilus subunit-phospholipid (VirB2-PG) unit of R388 polymerises into a 5-start helical filament in a manner similar to that observed in other conjugative pili (see strands named +2 to -2 in Figure 5A; hereafter, we refer to each unit as x_y_, where “x” is from a to c with subunits a to c being adjacent in the same strand; in this notation, “a” is under “b” and “c” is above “b”; subscript “y” indicating the strand in which the subunit is located). Both the rise and twist helical parameters are very similar among all conjugative pili (Figure S2). For R388, these parameters are 13.2 Å and 28.9°. In R388, this assembly results in a pilus with an interior lumen of 26 Å and external width of 82 Å (Figures 5A and S2), similar to other conjugative pili. As in pED208, all lipid headgroups in R388 are directed to the lumen (Figure 5B), changing dramatically the electrostatic potential of the lumen from negative to neutral (Figure 5B). Finally, while in pED208, each pilus subunit makes contact with 6 neighbouring subunits and 4 PG molecules, in R388 two more VirB2 subunits (8 total) are observed interacting with each other (Figure 6A-D). By doing so, overall buried surface area upon assembly accounts for 58 % of the protein surface and 93.5 % of the lipid surface, compared to 52 % and 87.5 % for pED208 for example (see details in Figure 6D).

**Figure 5.**
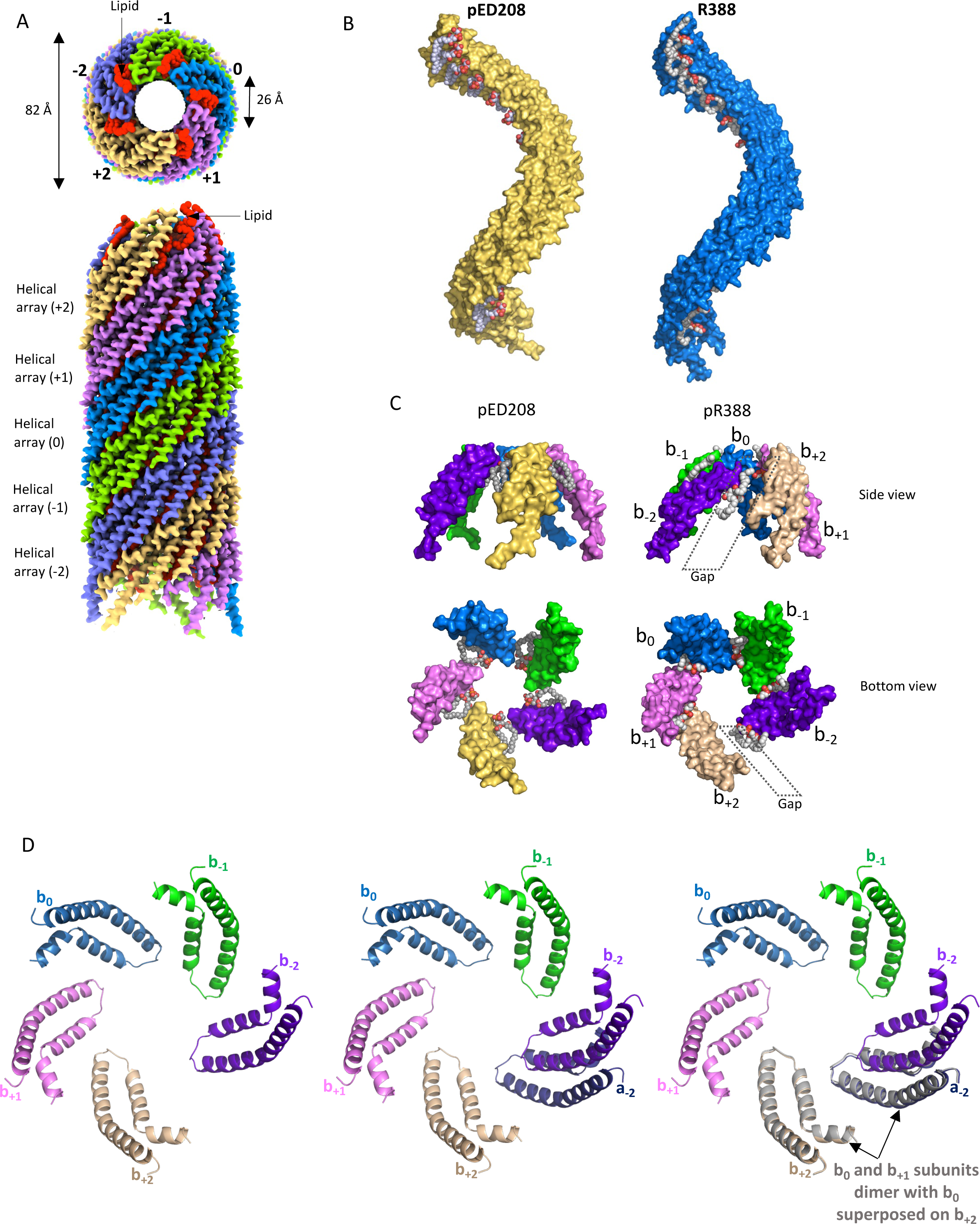
Helical assembly of the R388 pilus. (A). Top (upper panel) and side (lower panel) views of the R388 pilus. The structure is in surface representation. It consists of a 5-start helical assembly. Each of the 5 helical strands is shown in a different colour and is labelled -2 to +2. The colour coding for subunit in each strand is kept throughout the paper. (B). Comparison of the helical strands of pED208 and R388. The helical ladder of phospholipid along the strand is clearly visible in both. The VirB2 subunits are shown in surface representation while the lipids are shown in spheres representation colour coded by atoms type (grey and red for carbon and phosphorus/oxygen, respectively). (C). The pentameric layer of R388 (right) and comparison with that of pED2008 (left). Top panels: side view of both. Bottom: bottom view of both. Colour-coding and representation is as in B. Naming of subunits is introduced here and will be kept throughout the manuscript. Each subunit within a pentamer is ascribed a letter (b in this panel) and a subscript. The subscript refers to the strand from which the subunits shown are from (see panel A). the letter b is used here so that we can use in subsequent figures the letter “a” and “c” to indicate subunits of the pentamer located above or underneath the “b” pentamer, respectively. The gap between subunit b_+2_ and b_-2_ is shown. (D). The R388 pentamer is itself helical with a rise of 2.63 Å and a twist of 66.17°. Left: the R388 pentamer. Pentamer “b” is represented with colour coding of subunits as in panels A and C and naming b_-2_ to b_+2_. Proteins are shown in ribbon representation. Middle: pentamer “b” plus subunit a_-2_. Subunits a_-2_ and b_-2_ belong to the same helical strand. Right: a dimer made of subunits b_0_-b_+1_ in grey ribbon is used to superimpose the b_0_ subunit of this dimer onto b_+2_. This superposition results in b_+1_ of the grey dimer superimposing perfectly onto b_-2_, demonstrating the helical nature of the R388 pentamer.

**Figure 6.**
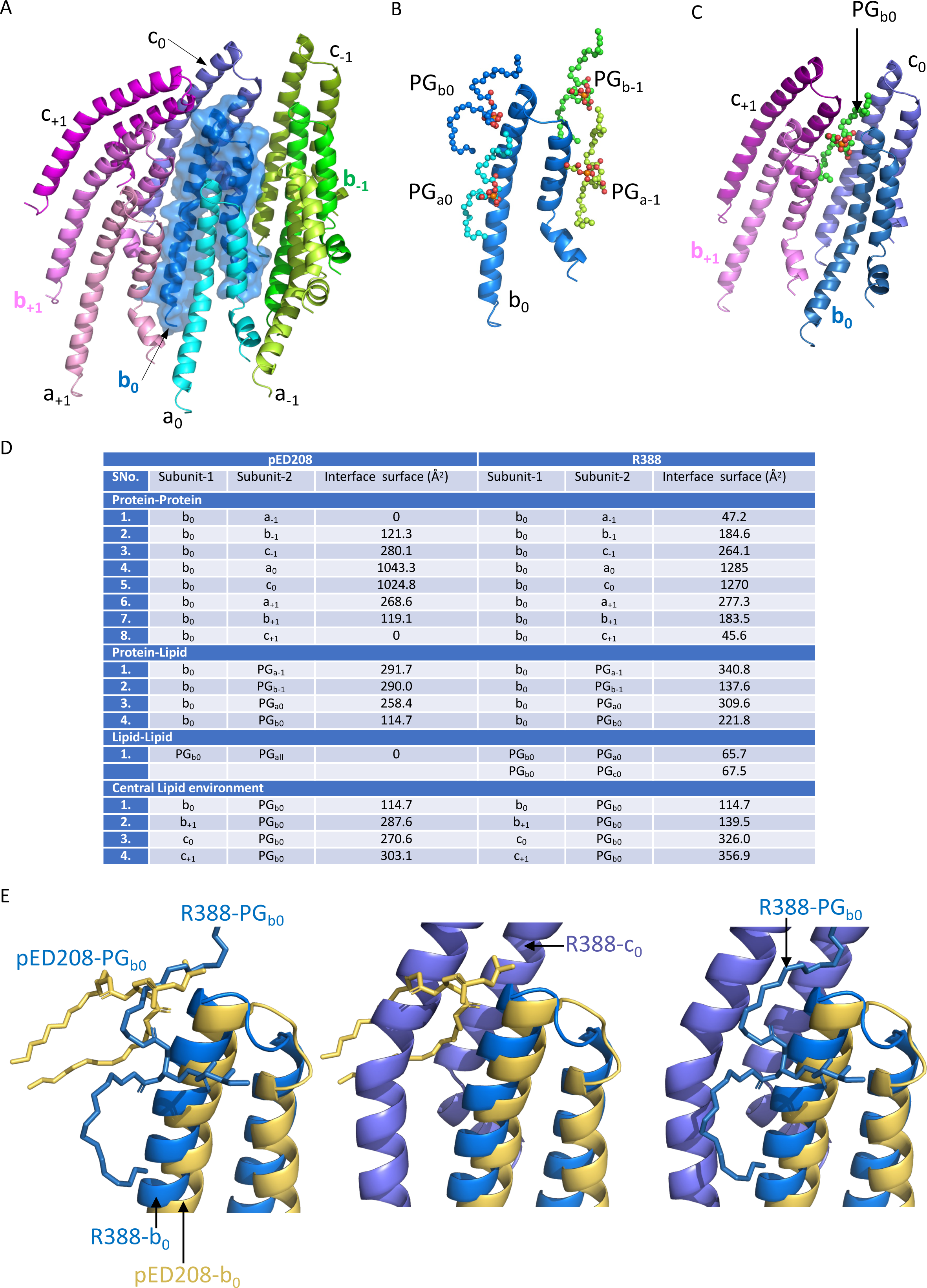
Protein-protein and protein-lipid interaction network. (A). Protein environment of the R388 b_0_ subunit. b_0_ makes contact with 8 adjacent subunits, two in the same helical strand (strand 0) and six in the immediately adjacent strands (strands +1 and -1). As mentioned in Figure 5C, “a” and “c” refer to subunits in the pentamers underneath or above the “b” pentamer, respectively. (B). Each subunit interacts with 4 lipids. All lipids are labelled according to the subunit they are bound to. For example, PG_b0_ represents the PG bound to b_0_. (C). Each PG makes interactions with 4 surrounding subunits. Here, PGb_0_ is shown surrounded by subunits b_0_, c_0_, b_+1_. c_+1_. (D). Buried surface area of all protein-protein, protein-lipid and lipid-lipid interfaces in both pED208 and R388. Notation for subunits in R388 and pED208 are the same. (E). The R388 PG is unable to adopt the same conformation as the pED208 PG. Left: the pED208-lipid (yellow) and R388-lipid (blue) complexes are superimposed together using the proteins as a guide. Proteins and lipids are shown in ribbon and stick representations, respectively. Middle: the pED208 and R388 are shown superimposed as in left panel. The pED208 lipid in stick representation is shown as is the subunit c_0_ (which is the subunit in the same strand as b_0_ but in the pentamer above b_0_). As can be seen, c_0_ heavily clashes with the pED208 acyl chains. Hence the need for *sn*-1 and *sn*-2 to splay in the R388 pilus (right panel).

Major differences between R388 and pED208 pilus assembly (and all other conjugative pili) are however evident. Previously, for pED208 and F, we showed that the VirB2-lipid polymerising units form successive horizontal layers of 5 units (Figure 5C), each stacking on top of each other according to the helical parameters specific for each pilus, resulting in the 5 helical strands filament shown in Figure 5A. Upon closer examination of one of these horizontal layers, depicted in Figure 5C, two notable differences are observed. Firstly, in pED208, the interaction between adjacent VirB2-PG units does not involve protein-protein contacts: they are uniquely mediated via the phospholipid (Figures 5C and 6D). This is not the case for R388 (Figures 5C and 6D). In R388, the interface has a large part involving lipids (360 Å^2^), but also protein-protein contacts are observed (47 Å^2^; Figure 6D and see details of interaction in “interaction between VirB2 subunits” section below).

Secondly, while the 5 units of the pED208 are 5-fold symmetrical, those of R388 are not. Indeed, as shown in Figure 5, C and D, the ring formed by the 5 units (b_-2_ to b_+2_ in the Figure 5, C and D; see explanation of subunit naming convention above and in figure legend) is open between two units (units b_+2_ and b_-2_ in Figure 5, C and D). The corresponding gap is about 8.4 Å wide at its narrowest point and 17.9 Å at its broadest point. We also observed a slight protrusion of the subunits relative to each other, suggesting that within the pentamer, subunits are related by helical symmetry. As demonstrated in Figure 5D, indeed, each polymerizing units in the pilus is related to the adjacent one by a twist angle of 66.2° and a small rise of 2.6 Å, and this applies along the entire pilus. Thus, two helical arrangements are apparent in the structure of the R388 pilus, but not in that of pED208. For clarity, in both sections below, we will refer principally to interactions as being either between helical strands (inter-strands) or within strands (intra-strands) where strands refer to the helical strands of the 5-stranded pilus filament as defined at the beginning of this section and in Figure 5, A and B.

### Interactions between VirB2 subunits

As mentioned above, each R388 VirB2 subunits interacts with surrounding 8 subunits, one on each side within the helical strand to which the subunit belongs (see subunits in hues of blue in strand 0 in Figure 6A) and 3 on each of the 2 strands adjacent (+1 in hues of magenta, and -1 in hues of green in Figure 6A). However, because of the helical symmetry within both vertical helical strands and pentameric base layers, there are only 4 unique interfaces which, ranked based upon surface buried area, are: i-the major one between adjacent subunits within helical strands (intra-strand; symmetry-related b_0_-a_0_ or b_0_-c_0_ in Figure 6, A and D; ∼1270 Å^2^), ii-two interfaces between subunits across the helical strands (inter-strand; symmetry-related b_0_-c_-1_ and b_0_-a_+1_ in Figure 6, A and D; ∼280 Å^2^; and symmetry-related b_0_-a_-1_ and b_0_-c_+1_ in Figure 6, A and D; 190 Å^2^); iii-a fourth between adjacent subunits within the pentameric base (also inter-strand; b_0_-b_-1_ and b_0_-b_+1_ in Figure 6, A and D; ∼50 Å^2^). When comparing to pED208 (or other conjugative pili) with the R388 pilus, there are two differences: the major interface of R388 buries 250 Å^2^ less than in pED208, and the b_0_-a_-1_ and b_0_-c_+1_ interfaces are not present in pED208, that interface being entirely mediated by the phospholipid in pED208 (Figure 6D) and other known conjugative pili.

The major interface, between subunits of the same helical strand, is primarily hydrophobic. It is too large to be described in detail and is similar to other intra-strands interfaces in other conjugative pili. However, the three other interfaces are much smaller and are described in detail in Figure S3. One notable feature is the fairly large number of residues of b_0_ interacting with both a_+1_ and b_+1_ (Figure S3), a reflection of the tight packing of the subunits in those regions.

### Protein-lipid and lipid-lipid interactions

As mentioned above, each VirB2 subunit interact with 4 phospholipids, and each phospholipid interacts with 4 VirB2 subunits. Overall, 1009 Å^2^ of surface area in each subunit is buried by the 4 PGs that interact with it, or 13.6 % of the total subunit’s solvent accessible area. Conversely, 937 A^2^ of surface area is buried in each PG upon interaction with the 4 surrounding subunits, or 76.7 % of the lipid’s solvent accessible area. Surface contact areas also vary from 115 Å^2^ for interaction of PG_b0_ with b_0_ to 360 Å^2^ for PG_b0_ interaction with subunit c_+1_ (Figure 6B; see details below). Finally, the PG of one subunit in one helical strand interacts with the PGs immediately above and immediately underneath in the same strand (66 Å^2^; interface PG_b0_-PG_a0_ or PG_b-1_ and PG_a-1_ in Figures 6B and 6C).

As previously highlighted, the conformation of the PG in R388 significantly deviates from that observed in other pili structures. While the acyl chains in all other pili typically exhibit parallel alignment, in R388, these chains assume a splayed configuration. The reason for this is apparent from comparing the packing of VirB2 pilus subunits in R388 and pED208 (Figure 6E). In Figure 6E, at left, we have superimposed the polymerising units of pED208 (in yellow) and R388 (in blue) using the two proteins as a guide (the two superimpose well with an RMSD of 2.21 Å in Cα positions). In the middle panel, we display in addition the intra-strand subunit of the R388 VirB2-PG unit immediately above in strand 0 (subunit labelled C_0_). This visualization reveals a substantial clash between subunit C_0_ and the acyl chains of the pED208 PG, necessitating the splaying of the PG acyl chains in R388.

In order to describe the details of protein-lipid interactions, we opted to focus on the 4 interfaces that 4 subunits make with one PG (as shown in Figure 6C), rather than focusing on the interactions that 4 PGs make with one protein (as shown in Figure 6B). These details are reported in Figure S4. Here we will focus on the main points that can be extracted from such details. Firstly, very few interactions are observed with the head group except for Y80b_+1_ and K87b_+1_ (sn3 in Figure S1; interaction details shown in Figure S4). Instead, a large number of hydrophobic residues from surrounding subunits appear to wrap around *sn*-1 and *sn*-2, contributing most of the binding contacts. Secondly, most of the interactions that *sn*-1 makes are with c_0_ with some minor contributions from b_0_ and b_+1_, while most of the interactions with *sn*-2 are contributed almost equally by a_0_, b_+1_, c_+1_, and b_0_.

### Validation of the structure

In order to examine the effect of targeted mutations, an assay for pilus biogenesis needed to be set up. Typically, this assay is based on imaging pili fluorescently labelled using maleimide coupled dyes reacting to a Cys introduced within the VirB2 pilus subunit at a position that does not affect function (Ellison et al., 2019). To that effect, we mutated independently four surface-exposed residues (hence likely to have no deleterious effect on function) to Cys and selected one, T63S, that offers best labelling efficiency and minimal effect on conjugation (Figure 7). With this pilus biogenesis assay in hand for the R388 system, we set out to observe the fluorescence patterns of live cells with or without recipient cells.

**Figure 7.**
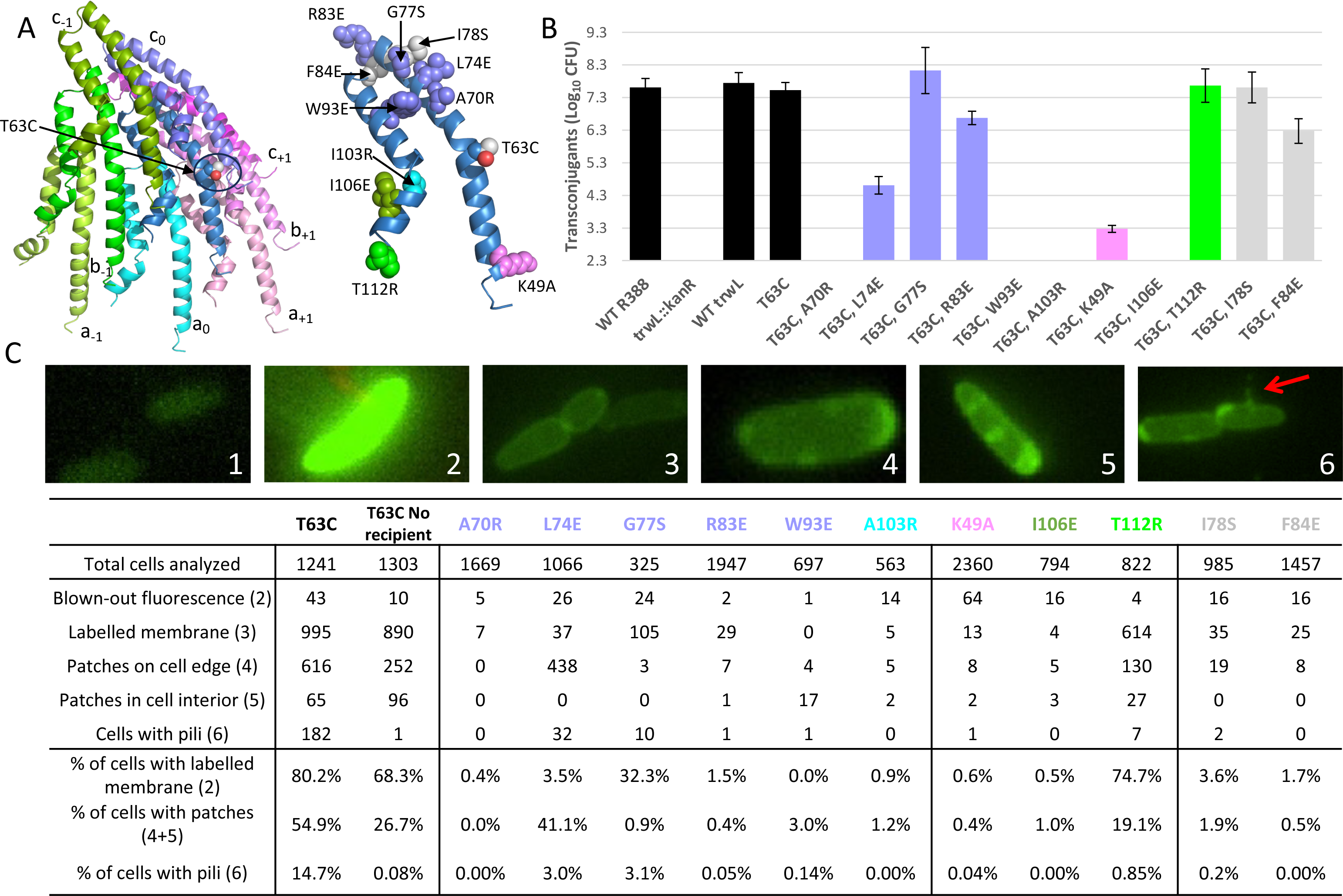
Point mutation analysis of TrwL/VirB2. (A). Locations of mutations made in this study. The orientation shown here is rotated by 180 degrees compared to what is shown in Figure 6A. Left: position of T63, the residue mutated to Cys and used to monitor fluorescently pilus biogenesis *in vivo*. Right: point mutations predicted to disrupt intra-strand interaction (residues in slate blue (b_0_-c_0_ interactions) and cyan blue (b_0_-a_0_ interactions)), inter-strand interactions (violet (b_0_-b_+1_ interactions), split-pea green (b_0_-a_-1_ interactions), and green (b_0_-b_+1_ interactions)), and interaction between lipids (grey). (B) Impact on conjugation efficiency of the mutations in (A). Unaltered R388 (WT R388), R388 with disruption of *virB2*/*trwL* (trwL::kanR), *virB2/trwL* disruption complemented with wildtype *virB2/trwL* (WT virB2/trwL), *virB2/trwL* disruption complemented with *virB2/trwL* with T63C mutation, and *virB2/trwL* disruption complemented with *virB2/trwL* with T63C mutation and mutations of intra- and inter-strand interface residues as described in main text. The Y-axis is set to begin at the minimum detection limit for the assay. (C). Microscopy analysis of pilus formation by maleimide-stained VirB2/TrwL mutants. Donors with each mutant was mixed with sfCherry-labelled recipient cells, imaged, then tallied as being: unlabelled cells (1), blown-out fluorescence (2), labelled membrane (3), fluorescent patches on cell edge (4), fluorescent patches in cell interior (5), and cells with visible pili (6). Cells could be counted in more than one category, though categories (3) and (4) were mutually exclusive. Analysis was also performed on the T63C mutant in the absence of recipient cells.

In addition to a subset of cells that always remained unlabelled, we also observed over-labelled cells with blown-out fluorescence. We attribute these cells to experimental artifact and excluded them from our analysis. Among the remaining cells, we observed four different types of fluorescent features (Figure 7C). The first was a clear membrane label, which we attribute to pilin subunits residing in the cell membrane. The membrane labelling was completely absent from cells lacking VirB2/TrwL or R388 plasmid. The second and third feature types were fluorescent patches either on the cell edge or within the cell interior, respectively. Cells that only had fluorescence accumulation at their poles were not counted. These patches appear to be VirB2/TrwL accumulations at pilus biogenesis sites. Whether these patches are on the cell’s edge or on the cells interior it is likely due to the rotational orientation of the cell on the microscope slide. The final feature we observed were appendages that extended outside of the cell, which appear to be the VirB2/TrwL pili (Figure 7C-far right panel labelled as number 6). For each observed pilus-like appendage, to gauge pilus dynamics, we collected an additional time-lapse recording lasting anywhere from 1 to 20 minutes. Although we were able to observe a couple instances of apparent pilus extension, we did not observe any cases of pilus retraction.

As mentioned above, during our biochemical work aiming to purify R388 pili in sufficient quantities to solve their structure, we observed a dramatic effect of the presence of recipient cells on pilus production. Thus, using our newly designed *in vivo* pilus production fluorescent assay, we next tested the effect of recipient cells on pilus production by donor cells (Table in Figure 7C). We observed a clear effect of recipient cell on pilus production. Indeed, while in the absence of recipient cell, only 0.08% of donor cells produce a pilus, 14.7 % of the donor cells produce a pilus in the presence of recipient cells. Note that the absence or presence of recipient cells does not affect the pool of VirB2 pilus subunits in the membrane. R388 is the first plasmid system where such observations are made. Previous investigations have shown that F-family plasmids (such as pED2008) do not require recipient cells for pilus production and thus, it was thought that pilus production does not need to be stimulated by the presence of recipient cells. This is not the case here: R388 is the first of its type being described where the presence of recipient cells is shown to have such a stimulatory effect.

In order to validate the structure, we next introduced mutations at various positions deemed to be structurally important to maintain helical packing and lipid binding. This validation exercise does not seek to be exhaustive as many conjugative pili structures have already been solved and mutational studies for most of them have been presented. For helical packing, we targeted i-residues involved in maintaining the largest interface between subunits, that within helical strands (helical strand 0 in Figure 7A for example, residues mutated: A70R, L74E, G77S, R83E, W93E), ii-residues involved in interacting across helical strands (Figure 7A; residues mutated: K49A, I106E, T112R), and iii-residues involved in PG-binding (Figure 7A; I78S, F84E).

Mutants that have the most pronounced detrimental effect on pilus production are K49A, A70R, L74E, A103R, I106E, W93E, validating the fact that the most important contacts are within the intra-strand (A70R, L74E, A103R, W93R) or the inter-strand (K49A and I106E) interfaces. T112R and G77S do not affect pilus biogenesis: T112 is solvent accessible and therefore might be able to adopt alternative conformations where contacts are no longer made with this residue across protein-protein interfaces; G77S is a small perturbation which might not be sufficient to disrupt significantly the interface in which this residue participates. Concerning the residues involved in interacting with PG, we observe no effect in I78S and an over 1 log difference in pilus production relative to wild-type for F84E. Since F84E is responsible for splaying one of the acyl chains (*sn*-2; Figure S4B), it is not surprising that it should have a significant effect on pilus production. The effect is however less severe than mutations that affect protein-protein packing, which is to be expected since the mutation made at F84 was meant to somewhat disrupt only PG binding but not the overall packing arrangement in which the PG participates.

### Conclusions

Our knowledge of conjugative pili encoded by T4SS has greatly expanded since the first structure of the pED208 and F pili. However, remarkable conserved themes have emerged, the most important of them is that they are all made from binary protein-phospholipid polymerising units. That encoded by the R388 plasmid is no exception, except that a splayed conformation of its phospholipid is observed. The presence of phospholipid has been hypothesized to lower the energetic barrier for re-insertion of pilus subunit into the membrane during pilus retraction. For all pili where retraction has been observed, the lipid in the VirB2-lipid polymerising unit adopt a conformation where the two acyl chains run parallel to each other, the same conformation that lipid adopts in the membrane. However, in R388, the acyl chains are splayed and, therefore, would need to undergo a reverse conformational change from spayed to parallel if the R388 pilus were to be able of retraction, a conformational change likely to be energetically costly. It is therefore not surprising that we do not observe retraction events for the R388 pilus.

Our study also reports on a unique requirement for recipient cells to stimulate R388 pilus production by donor cells. The mechanism by which this may occur is unknown. It could be that donor-recipient cell contacts are required or that a yet-to-be-identified soluble extracellular factor/molecule produced by recipient cells plays a role. Understanding how pilus production is induced by recipient cells will constitute an avenue of research for years to come. Given the importance of some of these plasmids in spreading antibiotics resistance genes, and the crucial role the pilus plays in this process, the elucidation of the details of regulatory loops and pathways leading to pilus biogenesis might lead to the design of novel tools and molecules able to control pilus production and, by so doing, prevent antibiotic resistance genes from spreading among bacteria populations.

Importantly, to accomplish the long-term research objectives of our project of dissecting the mechanism of DNA transfer through a T4S system, the elucidation of the R388 pilus structure, along with the aforementioned *in-vivo* assay, provides a robust structural framework to carry out exhaustive mutagenesis and identify phenotypes where the T4S system is trapped in a transfer-impaired intermediate state. Insights derived from these structures might unveil the substrate-recruitment and translocation strategies implemented by the T4S system to execute conjugation and these insights may be leveraged for the development of interventions capable of impeding the dissemination of antibiotic resistance genes within bacterial populations.

## Star Methods

### Purification of R388 pili

The R388 pilus encoded by the wild type R388 plasmid (trimethoprim resistant) was purified from the surface of *E. coli* Type-1 pilli deficient HB101 cells (carbenicillin resistant). Initially, an approach akin to the one outlined for F and pED208 pilus (Costa et al., 2016) was employed for R388 pilus production but resulted in notably low yields. Subsequent optimization efforts revealed a unique requirement for the presence of recipient cells to be in close proximity with the donor cells to induce R388 pilus production, distinguishing it from other conjugative pili (Amro et al., 2023; Costa et al., 2016; Kreida et al., 2023; Zheng et al., 2020). Thus, the strategy described in (Costa et al., 2016) was modified such that both donors and recipients are grown separately till the log-phage, spun down, resuspended, mixed together and plated on solid agar support for pilus production. 6L of both donor (WT-R388 plasmid containing *E. coli* HB101) and recipient cells (*E. coli* HB101 without R388 plasmid) were grown till ∼0.6 O.D_600_ from overnight pre-inoculum in Luria-Bertani (LB) media at 37 °C with trimethoprim (10 μg/ml) +carbenicillin (100 μg/ml) and only-carbenicillin (100 μg/ml) respectively. The cells from both these cultures were then pelleted down at 5,000g for 15 mins and resuspended in 5 ml LB per flask, mixed together and 10 ml of this mixed culture was plated onto one large (25X25 cm) Luria-Bertani (LB) media containing plate (a total of 6 plates used) for 90 mins at 37 °C for R388 pili production. The cells were then gently collected from plates using the SSC buffer (15 mM sodium citrate pH 7.2, 150 mM NaCl) and pili was gently shaved off from the cells by treatment with chilled SSC buffer for 2 hr at 4 °C under stirring, followed by two rounds of centrifugations at 4,000g for 20 min. The pili present within the resulting supernatant were precipitated by adding 5% PEG 6,000 and 500 mM NaCl. After an overnight incubation with 5% PEG 6,000 and 500 mM NaCl at 4 °C, the resultant precipitate was collected by centrifugation at 25,500g for 30 mins. The recovered pili from the precipitate were resuspended in 40 ml of water, followed by a slow spin at 5000g for 20 minutes to eliminate insoluble particles. Subsequently, a second round of PEG precipitation was conducted under identical conditions (5% PEG 6,000, 500 mM NaCl) for 1 hr at 4 °C and the pellet collected after a spin of 27,000g for 30 mins was resuspended in 1 ml of PBS (pH 7.4) buffer. The suspension was layered onto pre-formed CsCl step gradients (0.25–1.5 g/cm^3^) in PBS and centrifuged at 192,000g for 17 hr at 4 °C. The gradient was fractionated and the fractions containing the R388-pili were further purified by undertaking a sucrose density gradient ultracentrifugation using a 15-60 % sucrose density gradient made in the same buffer and centrifuged at 99,223g (using a SW40Ti rotor) at 4 °C for 18 hrs. The fractionated samples from the gradient were assessed for the presence of R388-pilin (TrwL) using SDS-PAGE/negative stain EM and the fraction containing the sample was extensively dialyzed against PBS (pH 7.4) to remove any traces of sucrose. Purity of the R388-pilin (TrwL) samples was analysed by SDS-PAGE followed by silver staining and negative stain EM while the identity of TrwL was verified by LC-ESI MS/MS.

### Cryo-EM grid preparation and data collection

C-flat grids (Protochips, USA; 1.2/1.3 400 mesh) were negatively glow discharged using PELCO Easiglow (Ted Pella, USA) and coated with graphene oxide. 3 µl of the purified pili sample was applied on each grid and a Vitrobot Mark IV (Thermo Fisher Scientific, USA) operating at 4 °C and 100 % humidity was used to incubate the sample on the grid for 30 secs and blotting for 16 secs (blot force -10) prior to vitrification in liquid ethane. The R388 pilus data were collected at the ISMB Birkbeck EM facility using a Titan Krios microscope (Thermo Fisher Scientific, USA) operated at 300 keV and equipped with a BioQuantum energy filter (Gatan, USA) with a slit width of 20 eV. The images were collected with a post-GIF K3 direct electron detector (Gatan, USA) operating in super resolution mode, at a magnification of 81,000 corresponding to a pixel size of 1.067 Å. The dose rate was set to 14.62 *e* per pixel per second and a total dose of 34.67 *e* per Å^2^ was fractionated over 50 frames. Data were collected using the EPU software with a defocus range −0.9 μm - −2.4 μm and a total of 4884 movies were collected.

### Cryo-EM data processing

Raw movies were imported in Relion 4.0 and MOTIONCOR2 (Zheng et al., 2017) was used for motion-correction and dose weighting. Motion corrected micrographs were then imported into CRYOSPARC v4.3.1 (Punjani et al., 2020) and patch-CTF (multi) was used for determining the defocus values. A few micrographs were first picked manually and 2D classification of these particles resulted in some 2D-templates which were then used to pick particles from the entire dataset using Filament-Tracer (auto-picking job for filaments in CRYOSPARC). The resulting picks from the autopicking job were manually curated using the “inspect picks” feature in CRYOSPARC and a total of 1,158,759 particles were then extracted with a box size of 400 pix. After multiple rounds of 2D classifications, a total of 209,930 good particles were taken forward for 3D reconstruction. Initially, one round of 3D-refinement (called Helix-Refine for helical filaments in CRYOSPARC) was carried out in C1 symmetry with no helical parameters and this reconstruction was used as an input to determine the correct helical parameters for the R388 pilus. A broad range of helical parameters, namely helical-rise between Δz: 10-15 Å, and helical twist between Δϕ 20-45° was initially used to scan the correct symmetry of the R388 pilus using the Symmetry Search feature in CRYOSPARC. This resulted in the determination of Δz and Δϕ for the R388 pilus at Δz: 13.25 Å and twist Δϕ 29.12° respectively. Following a 3D-refinement (called Helix-Refine for helical filaments in CRYOSPARC) job in C1 symmetry using these helical parameters, another round of symmetry search scanning, but this time with a finer range of Δz values between 12-14 Å and Δϕ between 28-30° was carried our resulting in the determination of the refined helical parameters for the R388 pilus as Δz: 13.221 Å and Δϕ: 29.03°. 3D-refinement (called Helix-Refine for helical filaments in CRYOSPARC) using the helical parameters (Δz, Δϕ) of (13.221 Å, Δϕ: 29.03°) resulted in a map at an average resolution of 3.46 Å. Attempts to improve the resolution of the map by carrying out CTF-refinement on the clean particle set (of 209,930 particles) resulted in a final map with an average resolution of 3.35 Å (and a LocRes resolution of 3.39 Å) and this map was subsequently used for model building. The final values of the helical parameters used in reconstructing this map (at 3.35 Å) by the Helix-Refine job was determined to be: Δz: 13.241 Å and Δϕ: 28.983°. Determination of the correct helical parameters from the layer lines within the power-spectra using Helixplorer-1 (Estrozi et al., 2018) was also used to independently verify whether the values obtained through the Symmetry Search in CRYOSPARC are correct.

### Model building

Using the sharpened map of the R388 pilus reconstructed through the helical image processing pipeline in CRYOSPARC, model building for the protein-part was carried out using the automated model building methodologies implemented in ModelAngelo (Jamali et al., 2023). The protein sequence of the R388 pilus monomer: TrwL/VirB2 and the CRYOSPARC sharpened map of the R388 pilus were used as the input for ModelAngelo and multiple rounds of optimization generated a structure with all the protein-monomers built within the R388 pilus map. This model was subsequently improved by manually inspecting and adjusting the fit of the main-chain Cα and amino acid side chains for each of the monomers within the density in COOT v0.9.3 (Emsley and Cowtan, 2004). One important feature of this methodology is that every monomer within the pilus is considered an independent entity and not related to any helical symmetry or symmetric packing arrangements during model building. This methodology was important in allowing us to deal with the defect in packing between the helical filaments within the R388 pilus as described above. The protein-subunit built model of the R388 pilus clearly highlighted unaccounted densities closer to the lumen of the fibre. However, interpreting and building a lipid molecule within these densities was more challenging owing to the differences in location as well as conformation of the lipid (with respect to the protein-monomer) as compared to all other known conjugative pili structures. Since the lipidomic analysis revealed that the lipid bound to TrwL is PG 32:1, several attempts were made to place the lipid in conformations similar to the ones seen in other conjugative pili structures with acyl chains running parallel to each other and did not succeed owing to the clashes with respect to the protein chain of the neighbouring subunit. Thus, to fit the lipid, a bundle of 9 monomers, comprising of one central monomer and 8 neighbouring monomers surrounding it was selected towards the centre of the fibre. First, atoms corresponding to the glycero-3-phospho glycerol group were placed within the density adjacent to the central subunit and the corresponding acyl chains were extended on each side to complete the lipid molecule. Since the lipid from the neighboring subunits are also situated in close proximity with the lipid of the central monomer, attempts were made to first fit the lipid in all the monomers in the immediate vicinity of the lipid corresponding to the central monomer and its conformation was optimized manually in COOT to ameliorate any clashes with respect to the protein monomer as well as the lipids/monomers of the adjacent subunits. Once the conformation of the protein-subunit and the lipid within the central monomer was finalized, this basic assembly was refined in PHENIX v1.20.1 (Adams et al., 2010) generating a refined monomeric unit that was superimposed on all the subunits within the pilus leading to a model of the pilus which has lipid bound to all the protein subunits. This model was subsequently refined in PHENIX v1.20.1 and following manual inspection and readjustment of all the subunits in COOT was refined again in PHENIX v1.20.1 to generate the final model of the R388 pilus.

### Mass spectrometry analysis of lipids

Lipid extractions from purified pili were achieved by 3 successive vigorous extractions with ethanol (90% v/v) (Fyffe et al., 2006). The pooled extracts were dried by nitrogen gas in a glass vial and re-extracted using a modified Bligh and Dyer method (Richmond et al., 2010). For whole cell control, membranes were washed with PBS and extracted following the procedure. Pili were treated with Phospholipase A2 (0.1 units) in PBS for 16 h at 37 °C, followed by heat inactivation and extraction as described above. PLA2 treated whole cell extracts were treated in the same manner and extracted using the same protocol.

Extracts were dissolved in 15 μl of choloroform:methanol (1:2) and 15 μl of acetonitrile:propan-2-ol:water (6:7:2) and analyzed with both a Thermo Exactive Orbitrap mass spectrometer by direct infusion and a ABsceix 4000 QTrap, a triple quadrupole mass spectrometer equipped with a nano-electrospray source. Samples were delivered using a Nanomate interface in direct infusion mode (∼125 nl/min). Lipid extracts were analysed in both positive and negative ion modes using a capillary voltage of 1.25 kV. MS/MS scanning (daughter, precursor and neutral loss scans) were performed using nitrogen as the collision gas with collision energies between 35-90 V.

### Bacterial growth and mutant construction

Imaging of the pili was done in strain NEB10 (New England Biolabs). Media with antibiotics used to select different plasmid/genetic elements: NEB10 strain (streptomycin, 50 µg/ml), R388 (trimethoprim, 10 µg/ml), pBAD24/pKD46 plasmids (Carbenicillin, 100 µg/ml), *trwL* deletions (kanamycin, 50 µg/ml), and pBAD33 plasmids (chloramphenicol, 15 µg/ml).

To disrupt the native *trwL* gene in R388, λ-red recombineering (Datsenko and Wanner, 2000) was used. The kanamycin resistance gene from plasmid pCOLADuet-1 was amplified using primers SX42 and SX43 (Table S3) and transformed into NEB10 competent cells carrying R388 and λ-red plasmid pKD46. Cells were then grown at 42°C to cure the pKD46 plasmid.

To generate TrwL mutants, the *trwL* gene was amplified by PCR using primers listed in Table S3, then cloned into plasmid pBAD24 between *Eco*RI and *Hind*III restriction sites. This construct (pBAD24-trwL) was then transformed into NEB10 chemical competent cell. The pBAD24-trwL construct was used as the template for site directed mutagenesis using KLD enzyme mix (New England Biolabs). Mutations were confirmed by sequencing.

### Conjugation efficiency assay

The donor strain was generated by transforming each of the pBAD24-TrwL mutants into NEB10 cells carrying the R388 *trwL::kanR* plasmid. The *E. coli* HST08 carrying pBAD33 to confer chloramphenicol resistance was used as the recipient strain. Donor and recipient cells were each grown in LB medium at 37 °C overnight in a shaking incubator (200 rpm), then sub-cultured (100x dilution) and grown to mid-log phase (OD_600_ ≈ 0.5). Donor cells were sub-cultured with 0.02% arabinose to induce the expression of mutant TrwL. The mid-log phase donor and recipient cells were spun down (5,000g for 2 min) and resuspended to OD_600_=10.0. Concentrated donors and recipients were mixed at a 1:1 ratio. 5 µl of the mixture was then spotted onto an LB agar plate, incubated at 37 ℃. After 2 hrs, the spotted cultures were cut from the agar plate, resuspended in 1mL LB, serially diluted, and spotted onto LB agar plates containing trimethoprim and chloramphenicol to count transconjugant CFU.

### Fluorescence microscopy imaging

NEB10 cells with R388 *trwL::kanR* and pBAD24-TrwL mutants were stained by maleimide-fluorescence dye (DyLight 488, Thermo Scientific) for pili visualization similar to previously described (Ellison et al., 2019). Pili-producing bacteria were grown in M9 + 0.4% casamino acid minimal media supplemented with LB (15 g/L) and appropriate antibiotic at 37°C overnight in a shaking incubator (200 rpm), then sub-cultured in LB + M9 + 0.4% casamino acid] with 0.02% arabinose until mid-log phase (OD_600_ ≈ 0.5). 1 ml of these donor cells were resuspended in 200 µl and 1 µl of maleimide-Dylight 488 solution (final dye concentration of 25 µg/ml) at 37℃ in a non-shaking incubator for 30 min. After staining, cells were spun down at low-speed (2,000g, 5 min) and resuspended in 50 µl of M9 + 0.4% casamino acid minimal media. The stained donor cells were mixed with recipient NEB10 cells carrying pBAD33-sfCherry. 0.5 µl of this cell mixture was spotted onto a 1% agarose pad (M9 + 0.4% casamino acid) for fluorescence microscopy.

### Image processing and data analysis

The fluorescence images were captured using a Nikon ECLIPSE Ti2 inverted microscope with a CoolLED pE4000 illuminator, Zyla 4.2 Megapixel Camera, and NIS-elements software. The microscopy images were analysed by the image processing algorithm MicrobeJ (Ducret et al., 2016) and Fiji (Schindelin et al., 2012).

## Acknowledgements

This work was funded by Wellcome grant 217089 to GW. We thank Dr Natasha Lukoyanova for data collection on the Birkbeck in-house Krios.

## Authors contributions

AV and AR produced and purified the pili, prepared cryo-EM grids, processed the cryo-EM data, built the model, made some of the mutants, and made some of the figures. SL set up the pilus biogenesis visualisation assay, made the mutants, and collected fluorescent imaging data. BH supervised the *in-vivo* pilus biogenesis work. TS undertook the lipid extraction and analysis and made figure 2. GW supervised the cryo-EM work, wrote the paper and made some of the figures.

