## Supplemental Tables and Figures for "Cryo-EM Structure of the R388 plasmid conjugative pilus reveals a helical polymer characterised by an unusual pilin/phospholipid binary complex"

**Table S1**. Cryo-EM data collection and model refinement statistics.

| Complex/Subcomplex | Conjugative pilus from *E. coli* R388 plasmid |
|---|---|
| **EMDB entry** | EMD- 19758 |
| **Data collection and processing** | |
| Magnification | 162,000 |
| Voltage (kV) | 300 |
| Electron exposure (e⁻/Å²) | 34.67 |
| Defocus range (μm) | -0.9 to -2.4 |
| Pixel size (Å) | 1.067 |
| Data Processing type | Helical refinement |
| Symmetry imposed | C1 |
| Initial particle images (no.) | 1,158,759 |
| Final particle images (no.) | 209,930 |
| Map resolution (Å) | 3.39 |
| FSC threshold | 0.143 |
| Map resolution range (Å) | 3.39-4.0 |
| Map sharpening B factor (Å²) | -109.7 |
| **Model building and validation statistics** | |
| Part(s) built in the map | TrwL protein after processing (AA 44 to 112) |
| Modelling method | Modelangelo followed by Coot and Phenix |
| PDB entry | 8S6H |
| **Refinement** | |
| EMDB corresponding | EMD-19758 |
| Initial model used (PDB code) | *De-novo* built using Modelangelo |
| Model resolution (Å) | 3.7 |
| FSC threshold | 0.5 |
| CC Model vs. Data (mask) | 0.78 |
| **Model composition** | |
| Chains | 69 |
| Nonhydrogen atoms | 39192 |
| Protein residues | 4761 |
| Ligands | 69 (LHG) |
| **B factors (Å²)** | |
| Protein | 96.69 (100 – 30) |
| Ligand | 20 (20-20) |
| **R.m.s. deviations** | |
| Bond lengths (Å) | 0.003 |
| Bond angles (°) | 0.429 |
| **Validation** | |
| MolProbity score | 1.77 |
| Clashscore | 3.47 |
| Rotamer outliers (%) | 4.24 |
| **Ramachandran plot** | |
| Favored (%) | 97.12 |
| Allowed (%) | 2.88 |
| Disallowed (%) | 0 |

| | |
|---|---|
| **Lipid PDB code** | LHG |
| **Mean CC for ligands** | 0.63 |

**Table S2**. Difference in torsion angles corresponding to acyl chains *sn*-1 and *sn*-2 between pED208 and R388. See Figure S1 for definition of angles.

| **Name of the acyl chain torsion angle** | **Difference (Δ) (in degrees) pED208-R388** |
|---|---|
| β1 | 73.2 |
| β2 | -38.94 |
| β3 | 111.99 |
| β4 | -29.33 |
| β5 | 99.35 |
| β6 | 179.24 |
| β7 | -103.66 |
| β8 | -20.46 |
| β9 | -126.26 |
| β10 | -133.63 |
| β11 | -67.79 |
| β12 | 9.93 |
| β13 | -47.42 |
| β14 | 14.43 |
| β15 | -115.98 |
| β16 | 44.23 |
| γ1 | 35.24 |
| γ2 | 22.11 |
| γ3 | 13.9 |
| γ4 | -33.74 |
| γ5 | -119.67 |
| γ6 | 148.57 |
| γ7 | -64.74 |
| γ8 | -169.43 |
| γ9 | 14.47 |
| γ10 | -55.26 |
| γ11 | -122.47 |
| γ12 | 14.14 |
| γ13 | -164.75 |
| γ14 | 68.04 |
| γ15 | -115.16 |
| γ16 | -41.93 |

**Table S3.** Strains and primers used in this study

**Strains used in this study:**

| **Strains** | **Description** | **Source** |
|---|---|---|
| BHom158 | *E. coli* ,DH10β/ pCOLADuet | our laboratory's collection |
| Pilar4 | *E. coli* ,Top10/pKD46 | Generous gift from Dr Pilar Menéndez Gil |
| Pilar207 | *E. coli* ,NEB10/pBAD33_sfCherry | Generous gift from Dr Pilar Menéndez Gil |
| SXC71 | *E. coli* ,NEB10/ WT R388 | This study |
| SXE1 | *E. coli* ,NEB10/ KanR@trwL R388 | This study |
| SXE9 | *E. coli* ,NEB10/ KanR@trwL R388 +pBAD24_T62C trwL | This study |
| SXE10 | *E. coli* ,NEB10/ KanR@trwL R388 +pBAD24_T63C trwL | This study |
| SXE11 | *E. coli* ,NEB10/ KanR@trwL R388 +pBAD24_T107C trwL | This study |
| SXE12 | *E. coli* ,NEB10/ KanR@trwL R388 +pBAD24_S52C trwL | This study |
| SXE13 | *E. coli* ,NEB10/ KanR@trwL R388 +pBAD24_WT trwL | This study |
| SXE31 | *E. coli* ,NEB10/ KanR@trwL R388 +pBAD24_T63C, R83E trwL | This study |
| SXE32 | *E. coli* ,NEB10/ KanR@trwL R388 +pBAD24_T63C, K49A trwL | This study |
| SXE33 | *E. coli* ,NEB10/ KanR@trwL R388 +pBAD24_T63C, T112R trwL | This study |
| SXE34 | *E. coli* ,NEB10/ KanR@trwL R388 +pBAD24_T63C, A70R trwL | This study |
| SXE35 | *E. coli* ,NEB10/ KanR@trwL R388 +pBAD24_T63C, A103R trwL | This study |
| SXE36 | *E. coli* ,NEB10/ KanR@trwL R388 +pBAD24_T63C, I106E trwL | This study |
| SXE37 | *E. coli* ,NEB10/ KanR@trwL R388 +pBAD24_T63C, W93E trwL | This study |
| SXE39 | *E. coli* ,NEB10/ KanR@trwL R388 +pBAD24_T63C, F84E trwL | This study |
| SXE40 | *E. coli* ,NEB10/ KanR@trwL R388 +pBAD24_T63C, I78S trwL | This study |
| SXE41 | *E. coli* ,NEB10/ KanR@trwL R388 +pBAD24_T63C, G77S trwL | This study |
| SXE42 | *E. coli* ,NEB10/ KanR@trwL R388 +pBAD24_T63C, L74E trwL | This study |

**Primer used in this study:**

| **Primer** | **Sequence** | **Function** |
|---|---|---|
| SX21_R388_TrwM_EcoRI | ACTAGAATTCACCATGGCAACTTCAATCTTGACG | Introduce EcoRI and HindIII restrinction site to *TrwL* and Clone *TrwL* to pBAD24 |
| SX22_R388_TrwM_HindIII | GAGAAGCTTTTAGGTAAACAGCATGGCC | |
| SX23_R388_TrwM_S52C_F | GGTGTGTCCTCGAAACATTGCAGCA | *TrwL* S52C(AGC---TGT) |
| SX24_R388_TrwM_S52C_R | TCGCCTTTTCCAATCCTTGAG | |
| SX27_R388_TrwM_T62C_F | AACTTTGTACGATTGTGCCCATTGCG | *TrwL* T62C(ACG---TGT) |
| SX28_R388_TrwM_T62C_R | CCTGCTGCAATGTTTCGAGG | |

| | | |
|---|---|---|
| SX29_R388_TrwM _T63C_F | ACGTGTATTGTGCCCATTGCGG | *TrwL* T63C(ACG---TGT) |
| SX30_R388_TrwM _T63C_R | AAGTTCCTGCTGCAATGTTTC | |
| SX31_R388_TrwM _T107C_F | TTGTGCCATGCTGTTTACCTAAAAGC | *TrwL* T107C(ACG---TGT) |
| SX32_R388_TrwM _T107C_R | ATCTGGACGGCGGAAC | |
| SX42_kanR_pCOL ADuet_trwL_f | TCCTCGAAACATTGCAGCAGGAACTTACGACGATTGTGCCCAT TGCGGCCGCTGTCGCGCTAGCATGCCTATTTGT | Clone KanR sequence from pCOLADuet for TrwL disruption (λ-red system) |
| SX43_kanR_pCOL ADuet_trwL_r | GCGGAACCGGCAATGATGACGCCGATAGACCACCGCACAAAC GTATCCTTTTCTTAGAAAAACTCATCGAGCATCA | |
| pBAD24$_{T63C}$R83E trwL_F | CGCGGGCGAATTCATCGAAAAGGATACGTTTGTGCGGTGGTC | For generating R83E mutant on *TrwL* |
| pBAD24$_{T63C}$R83E trwL_R | ACAAACGTATCCTTTTCGATGAATTCGCCCGCGTATGCGATGC C | For generating R83E mutant on *TrwL* |
| pBAD24$_{T63C}$ K49A trwL_F | ATTGGAAGCGGCGAGGAGCGTCCTCGAAACATTGCA | For generating K449A mutant on *TrwL* |
| pBAD24$_{T63C}$ K49A trwL_R | CTCGCCGCTTCCAATCCTTGAGCCGCCGCC | For generating K449A mutant on *TrwL* |
| pBAD24$_{T63C}$ T112R trwL_F | GCTGTTTCGCTAAAAGCTTGGCTGTTTTGGCGGATGAGAG | For generating T112R mutant on *TrwL* |
| pBAD24$_{T63C}$ T112R trwL_R | TTTTAGCGAAACAGCATGGCCGTAATCTGGACGGCGGAACC | For generating T112R mutant on *TrwL* |
| pBAD24$_{T63C}$ A70R trwL_F | TGCGGCCCGTGTCATCCTGCTCTGCCTCGGCA | For generating A70R mutant on *TrwL* |
| pBAD24$_{T63C}$ A70R trwL_R | GGCAGAGCAGGATGACACGGGCCGCAATGGGCACAATAC | For generating A70R mutant on *TrwL* |
| pBAD24$_{T63C}$ A103R trwL_F | CGGTTCCCGCGTCCAGATTACGGCCATGCTGTTTACCTA | For generating A103R mutant on *TrwL* |
| pBAD24$_{T63C}$ A103R trwL_R | AGCATGGCCGTAATCTGGACGCGGGAACCGGCAATGATGAC | For generating A103R mutant on *TrwL* |
| pBAD24$_{T63C}$ I106E trwL_F | CGTCCAGGAAACGGCCATGCTGTTTACCTAAAAGCTTGGCTGT TTTG | For generating I106E mutant on *TrwL* |
| pBAD24$_{T63C}$ I106E trwL_R | GCCGTTTCCTGGACGGCGGAACCGGCAATGATGACG | For generating I106E mutant on *TrwL* |
| pBAD24$_{T63C}$ W93E trwL_F | TGTGCGGGAATCTATCGGCGTCATCATTGCCGGTTCC | For generating W93E mutant on *TrwL* |
| pBAD24$_{T63C}$ W93E trwL_R | GGCAATGATGACGCCGATAGATTCCCGCACAAACGTATCCTTT TCG | For generating W93E mutant on *TrwL* |
| pBAD24$_{T63C}$ F84E trwL_F | GGGCCGAGAAATCGAAAAGGATACGTTTGTGCGGTGGTCTAT CG | For generating F84E mutant on *TrwL* |
| pBAD24$_{T63C}$ F84E trwL_R | CACAAACGTATCCTTTTCGATTTCTCGGCCCGCGTATGCGAT | For generating F84E mutant on *TrwL* |
| pBAD24$_{T63C}$ I78S trwL_F | CCTCGGCAGCGCATACGCGGGCCGATTCATC | For generating I78S mutant on *TrwL* |
| pBAD24$_{T63C}$ I78S trwL_R | GCGTATGCGCTGCCGAGGCAGAGCAGGAT | For generating I78S mutant on *TrwL* |
| pBAD24$_{T63C}$ G77S trwL_F | CTGCCTCAGCATCGCATACGCGGGCCGATTCATCG | For generating G77S mutant on *TrwL* |
| pBAD24$_{T63C}$ G77S trwL_R | CGCGTATGCGATGCTGAGGCAGAGCAGGATGACAGC | For generating G77S mutant on *TrwL* |
| pBAD24$_{T63C}$ L74E trwL_F | CATCCTGGAATGCCTCGGCATCGCATACGCG | For generating L74E mutant on *TrwL* |
| pBAD24$_{T63C}$ L74E trwL_R | GATGCCGAGGCATTCCAGGATGACAGCGGCCGC | For generating L74E mutant on *TrwL* |

### Supplementary Figure legends

**Figure S1**. **Structure of PG32 bound to pED208 (top) and R388 (bottom)**. Structures are shown in stick representation colour-coded by atom types, carbon in green, oxygen in red and phosphorus in orange. All atoms are labelled according to Marsh (2003) (Marsh, 2003). All dihedral angles are indicated and named according to Marsh (2003) (Marsh, 2003).

**Figure S2**. **Helical assemblies of eubacterial conjugative pili**. From top to bottom: Pilus name, resolution at which its structure was solved, rise and twist angle of helical assembly, side view of the pilus with one helical strand shown in grey and external dimension indicated, top view with lumen dimension indicated. All VirB2 subunits are shown in surface representation. All phospholipids are in surface representation, colour-coded in red.

**Figure S3**. **Protein-protein interactions between adjacent strands within the R388 helical assembly**. Only inter-helical strand interactions are shown. Left panel: overview of the 3 interfaces shown at right. Right top panel: residues involved in the $b_0$-$c_{+1}$ interface; Right middle: residues involved in the $b_0$-$b_{+1}$ interface; Right bottom: residues involved in the $b_0$-$a_{+1}$ interface. Residues are in stick representation colour-coded by subunit. Residue names are reported.

**Figure S4**. **Protein-lipid interaction details**.

(A). Overview of the interfaces between proteins and lipid depicted in panel B.

(B). Details of the residues involved in the interface between strand +1 subunits with $PG_{b0}$ (the lipid bound to the $b_0$ subunit). Proteins are in ribbon representation except for residues involved in interaction which are in shown in stick representation colour-coded according to the colour of the subunit they emanate from. Lipid is as in Figures 6B or S1.

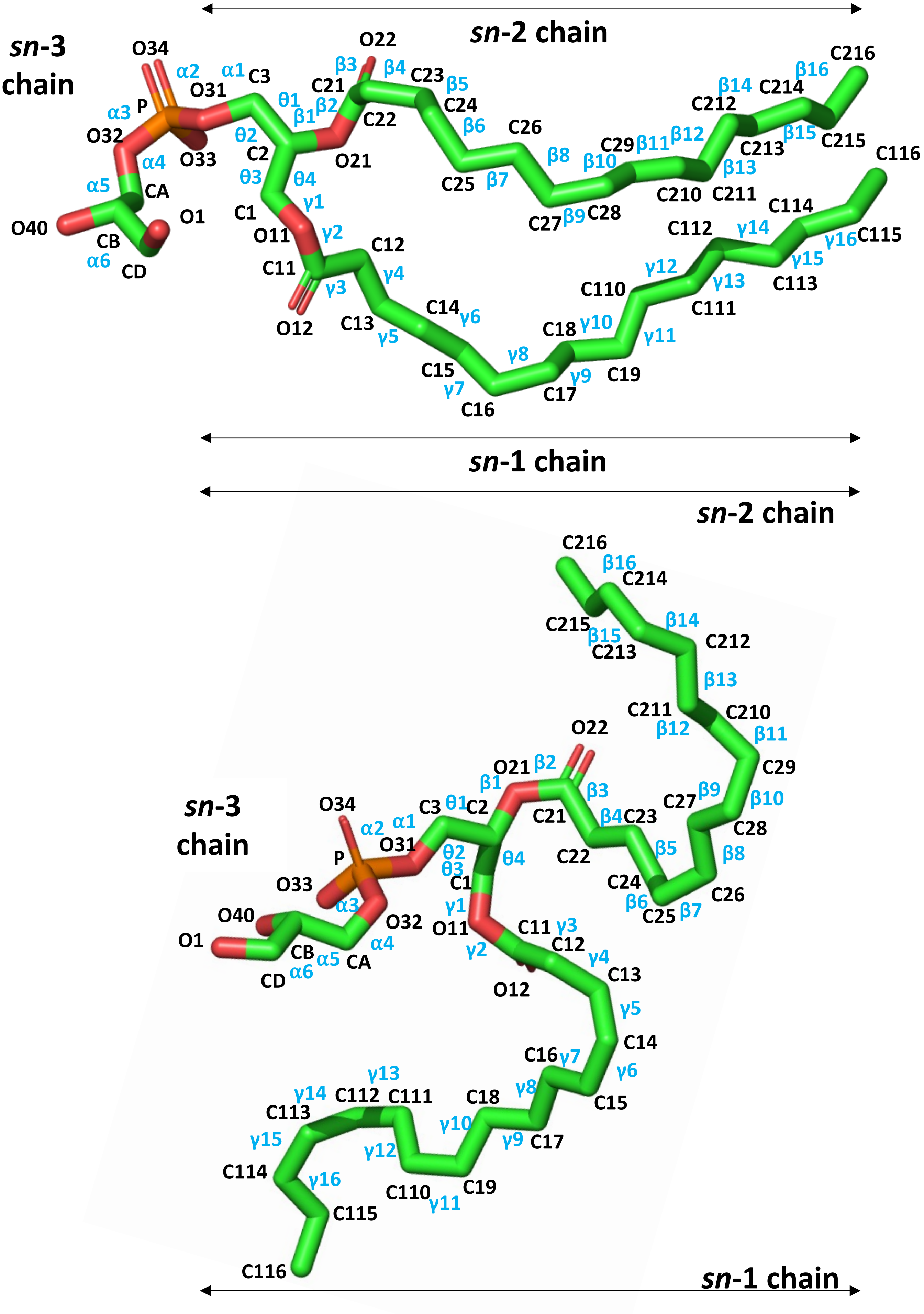
sn-2 chain
sn-3 chain
O34 α2 α1 C3
O31
α3 P
O32
O33
α4 CA
α5
O40 CB
O1
α6 CD
O22 β3 β4 C23 β5
C21 β2 C22
θ1 β1
θ2 C2 O21
θ3 θ4
C1 γ1
O11 γ2
C11 C12
γ3 γ4
O12 C13
C24 β6 C26
C25 β7 β8
C27 β9 C28
C29 β10 β11 β12
C210 C211 β13
C212 β14 C214 β16 C216
C213 β15 C215
C14 γ6 γ5
C15 γ7 C16
γ8 C17 γ9
C18 γ10 C19 γ11
C110 γ12 C111 γ13
C112 γ14 C114
C113 γ15 γ16 C115
C116
sn-1 chain
sn-2 chain
sn-3 chain
sn-1 chain

| | pED208 | F-pilus | pKpQIL | T-pilus | pKM101-pilus | R388-pilus |
|---|---|---|---|---|---|---|
| **Resolution (in Å)** | **3.6** | **5.0** | **3.90** | **3.50** | **3.00** | **3.20** |
| **Helical Parameters** | | | | | | |
| **Rise (in Å)** | **12.1** | **13.2** | **13.6** | **13.7** | **14.7** | **13.2** |
| **Twist (in °)** | **28.2** | **27.9** | **28.2** | **32.5** | **30.2** | **28.9** |

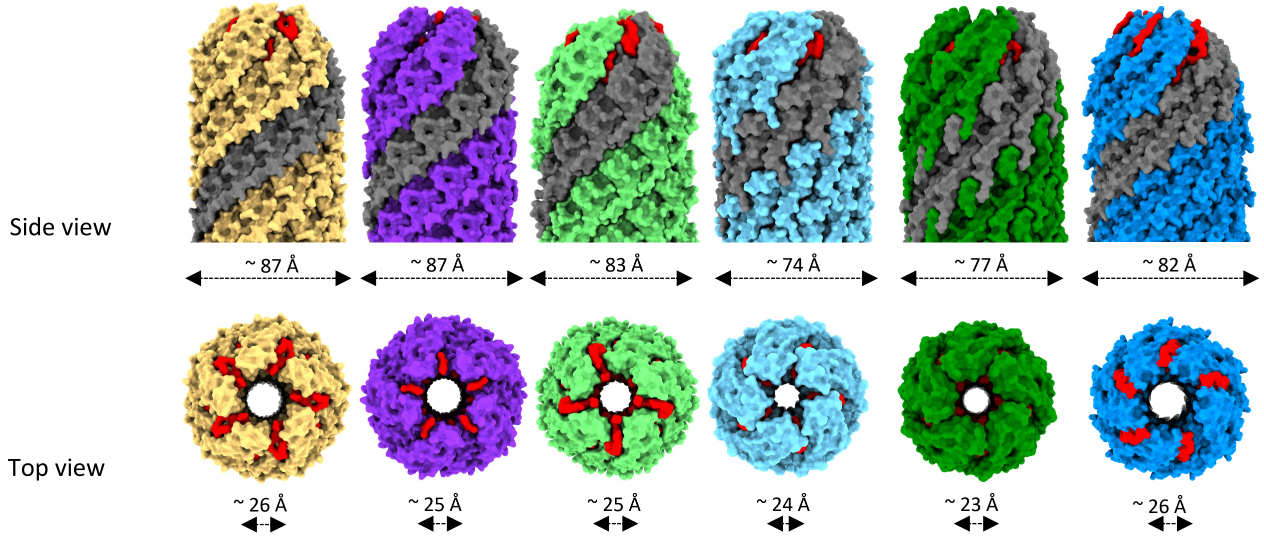

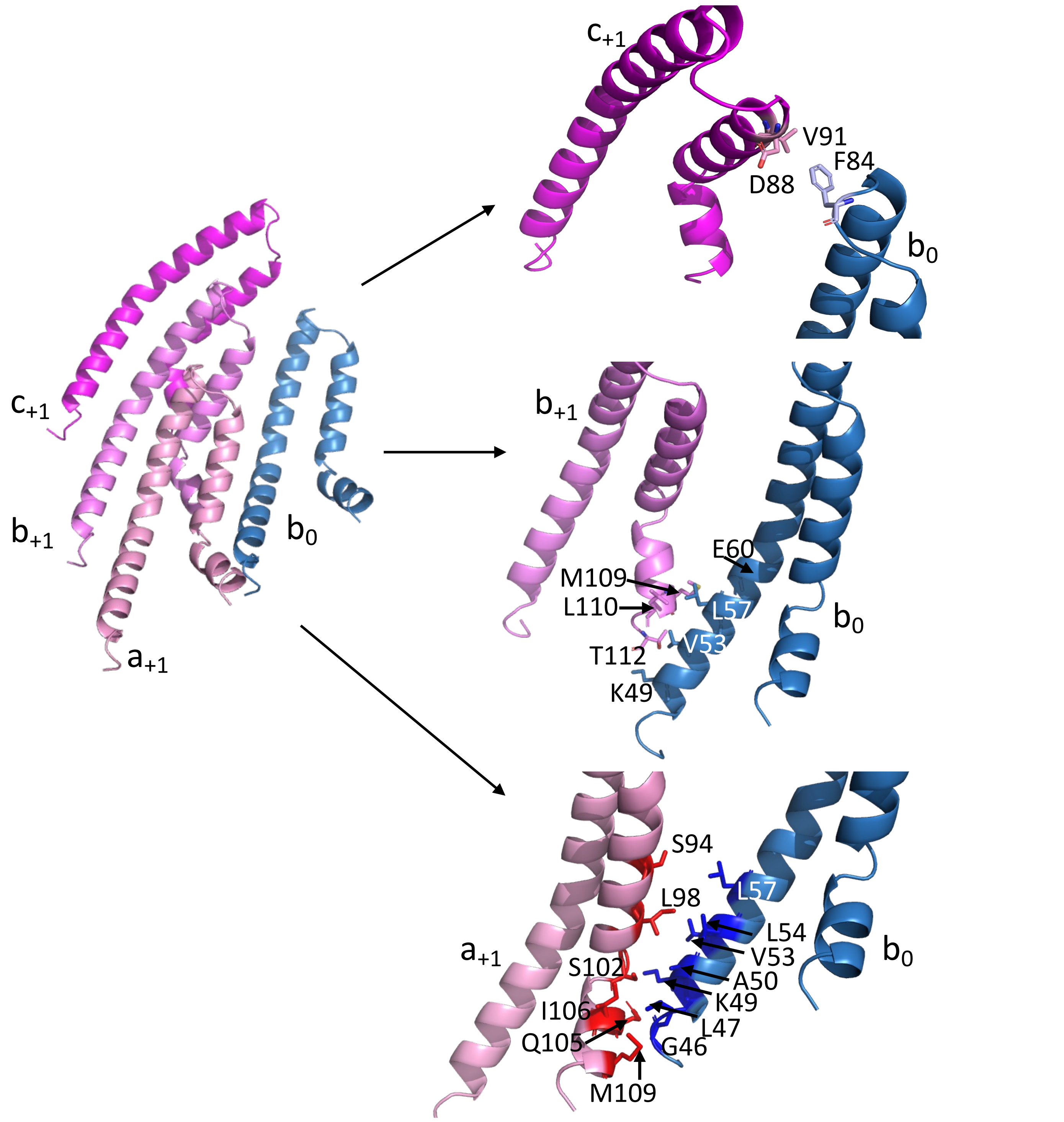

$c_{+1}$
$b_{+1}$
$a_{+1}$
$b_0$
$c_{+1}$
V91
F84
D88
$b_0$
$b_{+1}$
E60
M109
L110
L57
V53
T112
K49
$b_0$
S94
L57
L98
L54
$a_{+1}$
V53
S102
A50
K49
I106
L47
Q105
G46
M109
$b_0$

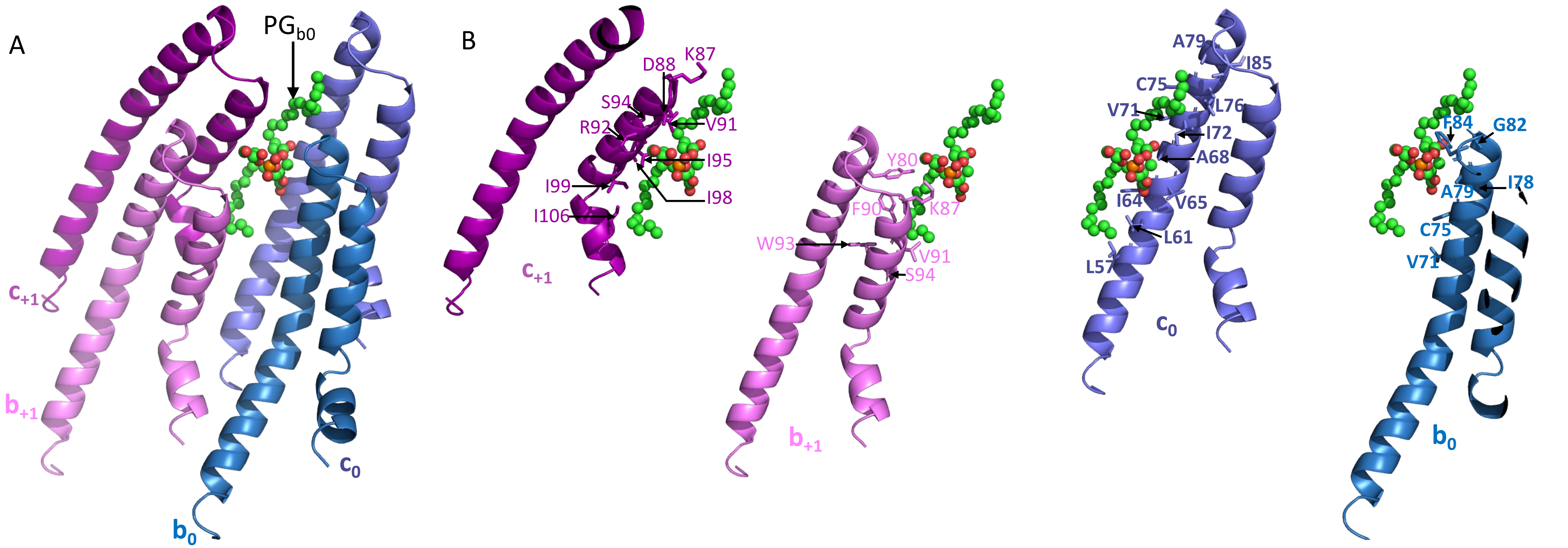
A
PG$_{b0}$
$c_{+1}$
$b_{+1}$
$b_0$
$c_0$
B
D88
K87
S94
R92
V91
I95
I99
I98
I106
$c_{+1}$
Y80
F90
K87
W93
V91
S94
$b_{+1}$
A79
I85
C75
V71
L76
I72
A68
I64
V65
L61
L57
$c_0$
F84
G82
I78
A79
C75
V71
$b_0$
